# Contextual enhancement on binge-like overconsumption of palatable sugar in mice

**DOI:** 10.1101/2024.09.29.615740

**Authors:** Hiroshi Matsui, Yasunobu Yasoshima

## Abstract

Binge eating disorder is an eating disorder characterized by the excessive intake of food within a short period, often beyond physiological needs. Studies using animal models have shown that binge eating animals consume food in quantities that surpass physiological necessity, and that the neural mechanisms underlying this behavior overlap with those involved in habit formation. Habitual behaviors are thought to be automatic responses acquired through extended behavioral training and are dependent on the context in which they were learned. Therefore, this study hypothesized that binge eating has a context-dependent component. We investigated whether the excessive palatable sugar intake observed in an animal model of binge eating is triggered by an associated context as a learned behavior. To test this, mice were trained to develop binge-like sugar consumption in a specific context. During the test phase, we reduced the animals’ hedonic needs for sugar solution by providing a two-hour satiety period. Sugar solution was then presented in both the training context and a novel context (Experiment 1) or a negative context (Experiment 4). The results showed that in the training context, the mice continued to consume sugar solution at levels similar to those observed at the onset of the satiation. In contrast, this context-induced sugar consumption was not observed in the novel context (Experiment 2), while a follow-up experiment employing conditioned place preference demonstrated the existence of contextual learning itself (Experiment 3). These findings collectively suggest that, like habitual behaviors, binge eating is induced under in the context-dependent manner and insensitive to the consequence of the behavior.

## Introduction

Binge eating disorder is a behavioral disorder characterized by the inability to control the consumption of large amounts of food in a short period (Brownley, Berkman, Sedway, Lohr & Bulik, 2007; Mathes, Brownley, Mo & Bulik, 2009). With a global prevalence ranging from 0.3% to 1.8% in adults, this disorder persists even after physiological hunger has been satiated, leading to significant impairments in psychological well-being and quality of life (Giel et al., 2022). Furthermore, binge eating disorder is often associated with obesity and food addiction, exacerbating its impact on individual health (Cassin & von Ranson, 2007; de Zwaan, 2001). The environmental and behavioral factors, along with the neural mechanisms underlying this disruption of feeding regulation, are actively explored (Keski-Rahkonen, 2021).

To investigate the mechanisms underlying binge eating, rodent models have been proposed (Oswald, Murdaugh, King, & Boggiano, 2011; Rossetti, Spena, Halfon, & Boutrel, 2014; Yasoshima & Shimura, 2015). For instance, in the protocol proposed by Yasoshima & Shimura (2015), mice were induced to develop binge-like excessive sugar intake through a regimen of successive restricted feeding. In their study, the animals were allowed access to normal chow and palatable sugar (sucrose solution) for only 4 hours each day over a period of 10 days. This repeated restricted feeding led the animals to consume significantly larger quantities of the sugar in a short time than they would normally. Notably, their study also revealed that this feeding behavior persisted even after the animals’ physiological need for nutrition had been met, demonstrating that the model successfully mimics clinical characteristics associated with binge eating disorder. The restricted feeding of chow and palatable foods can lead to behavioral alterations, and this procedure has been used in other studies as a training protocol for binge-like eating (e.g., Awad et al., 2024; King et al., 2016).

The nature of binge eating, marked by unnecessary feeding behaviors, resembles habitual behavior (LeMon, Sisk, Klump & Johnson, 2019; Moore, Sabino, Koob & Cottone, 2019). In psychology of learning, the habits are understood as the acquired automaticity of behaviors (Ciria, Watson, Vadillo & Luque, 2021). In the literature of instrumental learning, habitual behavior is operationally defined as decrement of sensitivity to the outcome produced by behavior (e.g., foods obtained via lever press). The instrumental behavior is typically sensitive to the outcome. For example, the lever press response is reduced by the taste aversion learning or satiation by free access to the reinforcer (‘outcome devaluation’; Adams, 1982; Adams & Dickinson, 1981). This mode of behavior is called ‘goal-directed’ (Dickinson & Balleine, 1994). It has been considered that the goal-directed behavior is emitted based on the current value of the reinforcer to fulfill the specific needs. In contrast, habitual behavior develops via extensive repetition of instrumental behavior, leading to being insensitive to the outcome (Dickinson et al., 1985; Dickinson, Balleine, Watt, Gonzalez & Boakes, 1995).

There may be a similarity between habits and overeating in that both involve seemingly automated behaviors, which are insensitive to the immediate needs of animals. Indeed, Furlong and Balleine (2017) demonstrated that the dorsolateral striatum is involved in binge-like behavior, which has been known to play a role in habitual actions, revealing a common neurophysiological basis. The maladaptation of corticostriatal circuits were also found in human neuroimaging (Kessler, Hutson, Herman, & Potenza, 2016). These similarity between binge eating and habitual behavior lead to the hypothesis that if similar mechanisms underlie both habits and binge eating, the factors that control habitual behavior might also apply to overeating.

Once a behavior becomes habitual, it is not necessarily irreversible. Recent research has shown that habitual behavior can be sensitive to the context in which it was developed (Steinfeld & Bouton, 2020, 2021; Thrailkill & Bouton, 2015). For example, Steinfeld & Bouton (2021) trained rats to perform a goal-directed instrumental response in one context (context-A) and then extended the training in a different context (context-B) to develop a habit. Devaluation through taste aversion learning was applied in both contexts, followed by an extinction test. The results showed that the instrumental response remained intact in context-B but not in context-A, indicating that habitual behavior is expressed in the context where it was acquired. Given the parallels between overeating and habits, it is plausible that overeating could also be understood as a context-dependent learned behavior.

The present study, therefore, investigated whether context could facilitate overconsumption by combining appetitive contextual conditioning with restricted feeding training to induce this behavior. The study consisted of two experiments: Experiment 1 examined whether binge-like overconsumption would be facilitated by the trained contexts after sensory-specific satiation. Experiment 2 assessed whether animals that underwent contextual conditioning but did not receive binge eating training would not show overconsuming palatable sugar after satiation. However, in this experiment, the mice were not subjected to food deprivation, which may have diminished their motivation to associate the context with the food. Therefore, in Experiment 3, we conducted training identical to that in Experiment 2 and subsequently assessed conditioned place preference to determine whether the mice perceived the context as a positive stimulus. Experiment 4 replicated Experiment 1 using female mice with a modified procedure. In Experiment 1, a novel context was used as a comparison to the trained context; however, this may have reduced responses due to neophobia. To address this issue, Experiment 4 replaced the novel context with a negative context, with which food was not paired.

## Materials and Methods

Subjects and housing conditions: total 51 C57BL/6J mice (CREA Japan, Osaka) were utilized for this study, with mice assigned to each of the four experiments (17 in Experiment 1, 17 in Experiment 2, 9 in Experiment 3, 8 in Experiment 4). The male mice were used in Experiment 1-3 and female in Experiment 4. Mice were purchased at eight weeks old and acclimated to the housing room for one week. They were housed individually in Plexiglas cages maintained at 22 °C under a 12L/12D cycle, with lights on at 7:00. Prior to the experiments, the mice had ad libitum access to water and a standard chow diet. The standard chow consisted of a pelleted commercial rodent diet (MF, Oriental Yeast Co., Ltd.: 3.6 kcal/g), which was used throughout the experiments. At the onset of the experimental procedures, the animals were 9 weeks old. All experimental procedures were conducted during the light cycle. All animals were treated in accordance with the NIH Guidelines for the Care and Use of Laboratory Animals. The study’s rationale, design, and experimental protocols were approved by the Institutional Animal Care and Use Committee of the Graduate School of Human Sciences, Osaka University (approval number: R6-3-0).

### Experimental procedure

*Experiment 1*: the experiment consisted of training and test phases, and both performed in the home cage. The restricted feeding procedure was utilized for shaping binge-like overconsumption (Yasoshima & Shimura, 2015; Figure 1a, top). All animals were maintained on a 20-h food deprivation schedule followed by 4-h access to the sucrose solution, water, and chow starting at 09:00 h every morning (2-h after lights on). The two bottles were presented side-by-side (two-bottle method): one bottle for the sucrose solution (0.5M concentration) and another for water. The water bottle was provided to ensure that the mice’s consumption of the sucrose solution was driven by the sucrose’s incentive rather than merely by thirst.

**Figure 1.**
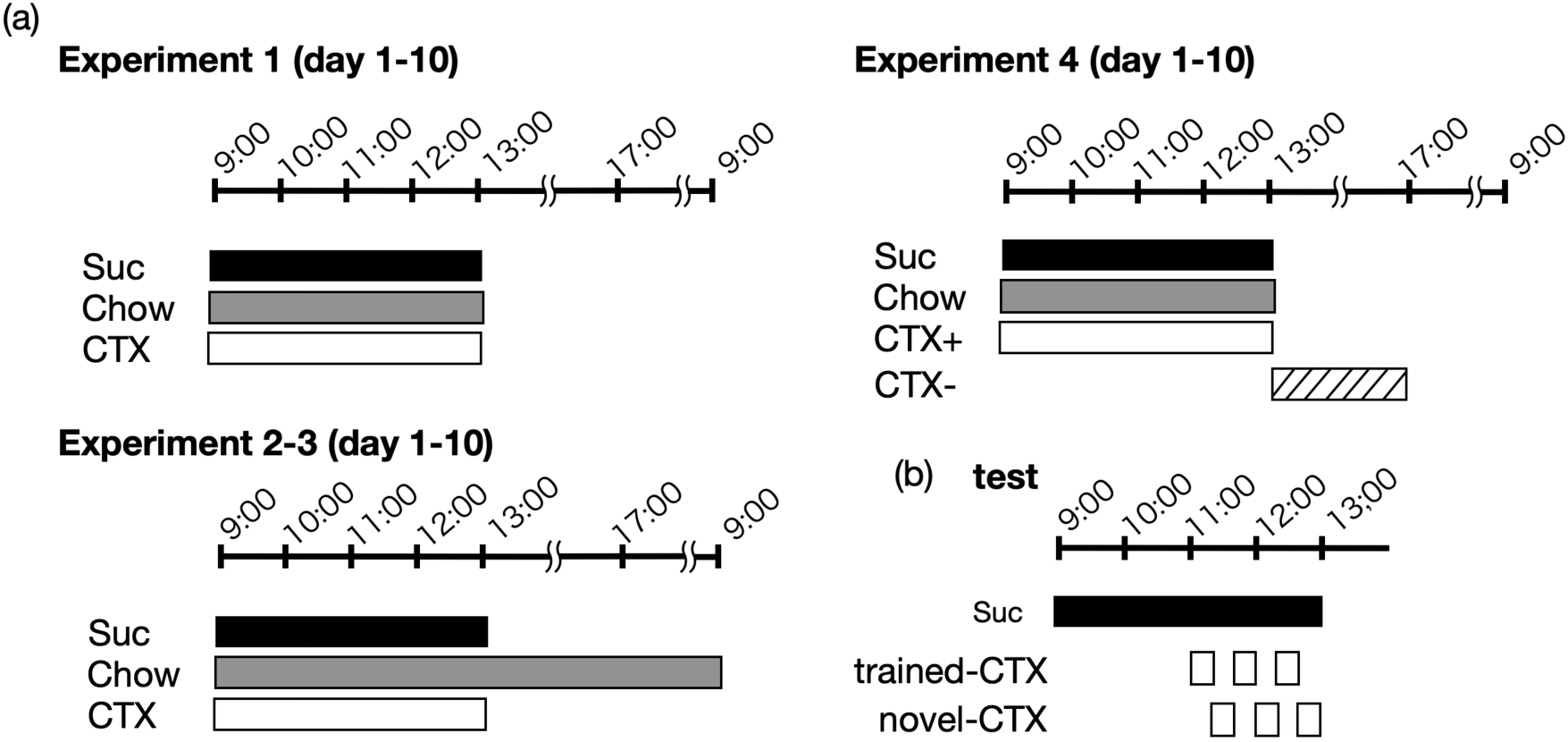
The timeline of the study included (a) a training phase and (b) a test. (a) Training was conducted over 10 consecutive days. (b) Testing took place immediately after the 10^th^ day of training. Half of the animals were first exposed to the trained context after a 2-hour period of satiation, while the other half were initially placed in a novel context. The illustration in the figure represents the time course of the group that started with the trained context. Suc: sucrose, Chow: standard chow, and CTX: context.

These 4-hour feeding occasions ran under the visual ‘context’. The context was as vertical or horizontal stripe walls made of acrylic plates. It was positioned to enclose all four sides of the home cage. Given that the context was always paired with palatable sugar and normal chows, the appetitive contextual conditioning was applied, where the context functions as the conditioned stimulus (CS) to signal the sugar/food presentation (US). The context was set just before beginning of the training, and immediately removed after 4-hour periods of food access. The vertical or horizontal patterns were balanced for arranging the groups in the test phase described below. The restricted-feeding training continued 10-days.

The test was conducted the day after the last training session (day-11; Figure 1b). Prior to testing, all animals were given 2-hour access to the sucrose solution to achieve sensory-specific satiation. The water bottle was likewise positioned in the same manner as during the training sessions. The satiation session was undertaken without the context. Consumption was measured at one-hour intervals. After the satiation, the animals were given the access to the sucrose solution under the context: trained or novel. The satiation procedure was performed to determine whether the mice would consume sucrose solution in excess of their hedonic needs within the trained context.

Thus, immediately after satiation, animals were again granted access to the sucrose solution for 20 minutes in either a trained or novel context. A novel context refers to one that was not used during training. For example, if an individual was trained in a context with vertical stripes, a novel context would involve horizontal stripes, and vice versa. We chose to use the novel context due to concerns that prior habituation to an untrained context might render it an inhibitory conditioned stimulus. Moreover, the facilitative effects of the trained context might not solely reflect the influence of the context itself, but could also be attributed to incidental cues, such as the experimenter’s handling of the context or the occlusion of the surrounding environment by enclosing walls. To eliminate these possibilities, it was necessary to use the novel context as a control condition rather than omitting the context presentation entirely. However, the novel context might induce neophobia, which could potentially interfere with sucrose intake. We aimed to rule out this possibility in Experiment 4.

After each feeding trial, the context was switched, allowing the animals another opportunity to feed. This procedure was repeated for three trials in each context, totaling six trials. Thus, the cumulative access time to the sucrose solution in each context was 1 hour. The animals were divided into two groups based on whether they started the test in the trained or novel context: the half of animals initially exposed to the trained context (i.e., trained -> novel -> trained -> novel -> trained -> novel), and another half of animals following in the reversed sequence.Group assignment was matched according to the sucrose solution intake on training day-10 to ensure no biases between the groups.

*Experiment 2*: most of the procedure was the same as Experiment 1 unless otherwise noted Figure 1a, bottom). The normal chow was ad libitum throughout the training periods unlike Experiment 1. The sucrose solution was presented in the same way as Experiment 1, from 9:00 to 13:00. Thus, the animals were exposed with paring of the context and sucrose solution. This protocol was adopted because Yasoshima & Shimura (2015) previously revealed that ad libitum to normal chow hindered the development of binge-like overconsumption. The test was performed completely the same as Experiment 1. This procedure allows for the contextual conditioning of sucrose solution consumption in animals without inducing binge-like eating behavior. The rationale for Experiment 2 is to determine whether sucrose solution consumption, triggered by the context, occurs after satiation as a result of the contextual conditioning experience alone.

*Experiment 3*: The training procedure was identical to that of Experiment 2. A potential issue in Experiment 2 was that the mice were not food-deprived, which may have resulted in insufficient motivation to consume the palatable sugar. Consequently, the association between the context and the food might not have been established.

To address this, in Experiment 3, instead of measuring food consumption in the context during the test, we assessed conditioned place preference (CPP) both before and after training. CPP was evaluated using the trained and novel contexts, as in Experiments 1 and 2. The experimental chamber consisted of two connected compartments, each with striped patterns on the walls corresponding to their respective contexts. The animals were placed in the chamber for 10 minutes and allowed to move freely between the two compartments. If contextual learning had occurred, even in the absence of food deprivation, the animals were expected to spend more time in the trained context during the post-test compared to the pre-test. The initial placement of the animals was counterbalanced across conditions.

*Experiment 4*: although we anticipated and, indeed, observed context-specificity in binge-like sugar consumption in Experiment 1, a major limitation was the exclusive use of male mice. Additionally, the novel context served as the reference for the trained context during the test session, which may have led to neophobia interfering with sugar consumption. To address these issues, Experiment 4 was conducted using female mice with a modified procedure based on Experiment 1. The training procedure was nearly identical to that of Experiment 1, where the training context was paired with food. The only difference was that another context was presented without food (Figure 1, top right). The exposure duration for this context was the same as that of the training context (4 hours). The testing procedure was identical to that in Experiments 1 and 2.

### Data Analysis

The increase in food intake during training was analyzed using one-way ANOVA with experimental sessions (days) as the independent variable. Satiation during the test was assessed by examining: 1) whether sucrose solution consumption decreased between the first and second hours, and 2) whether consumption differed from day 10 of training. A two-way ANOVA was conducted with day (day 10 and test) and time (the first and second 1-hour periods) as the independent variables. The Greenhouse-Geisser method was applied to adjust the degrees of freedom for within-subject factors. Additionally, the total consumption over these two hours was compared using a t-test.

Sucrose solution intake in the trained and novel contexts was compared using a t-test. For the trial-by-trial fluctuation of sucrose consumption during the test sessions, we adopted a linear mixed model to estimate the effect of trials and the context conditions, because the data is not complete; it is obvious that one animal could not simultaneously receive foods under different contexts (e.g., an animal which experienced the trained context in the first trial cannot receive the novel context in that trial). The conditions of contexts and trials were treated as fixed effects, and individual differences were accounted for as random effects.

For the CPP test in Experiment 3, we used markerless tracking software, DeepLabCut (Nath et al., 2019), to extract body center coordinates as the mice moved freely within the experimental chamber. The probability of staying in the trained context was quantified and compared between the pre- and post-tests using a t-test.

All analysis and visualization were performed under R version 4.1.0 using the packages including ‘lme4’ v1.1-27.1 and ‘car’ v3.1-2 for linear mixed model and the likelihood ratio test (lmer() and Anova() function), ‘rstatix’ v0.7.0 for ANOVA (anova_test() function), and ‘tidyverse’ v1.3.1for data handling and processing.

## Results & Discussion

### Experiment 1: contextual enhancement on the overconsumption of palatable sugar

The animals increased the feeding amount across the training periods, and reached the asymptotic levels in both normal chows and sucrose solution (Figure 2), replicating the previous binge-like consumption (Yasoshima & Shimura, 2015). Therefore, we consider that this restricted feeding regimen effectively worked as a model for binge-like eating. A one-way analysis of variance (ANOVA) revealed the significant effect of 1-hour consumption of sucrose solution (F(3.65,58.43) = 26.94, p < .0001), that of total 4-hours (F(4.35,69.52) = 120.79, p < .0001), 1-hour consumption of chow (F(4.51,72.20) = 12.57, p < .0001), and that of total 4-hours (F(2.95,47.16) = 28.39, p < .0001). On average, the animals consumed sucrose solutions more than 20% of animals’ body weights in the later sessions, whereas less than 10% in the initial sessions. The body weights also changed across training sessions ((F(1.26,20.22) = 39.96, p < .0001). The multiple comparisons revealed that body weights significantly increased after the 7-th session compared to sessions 1-7.

**Figure 2.**
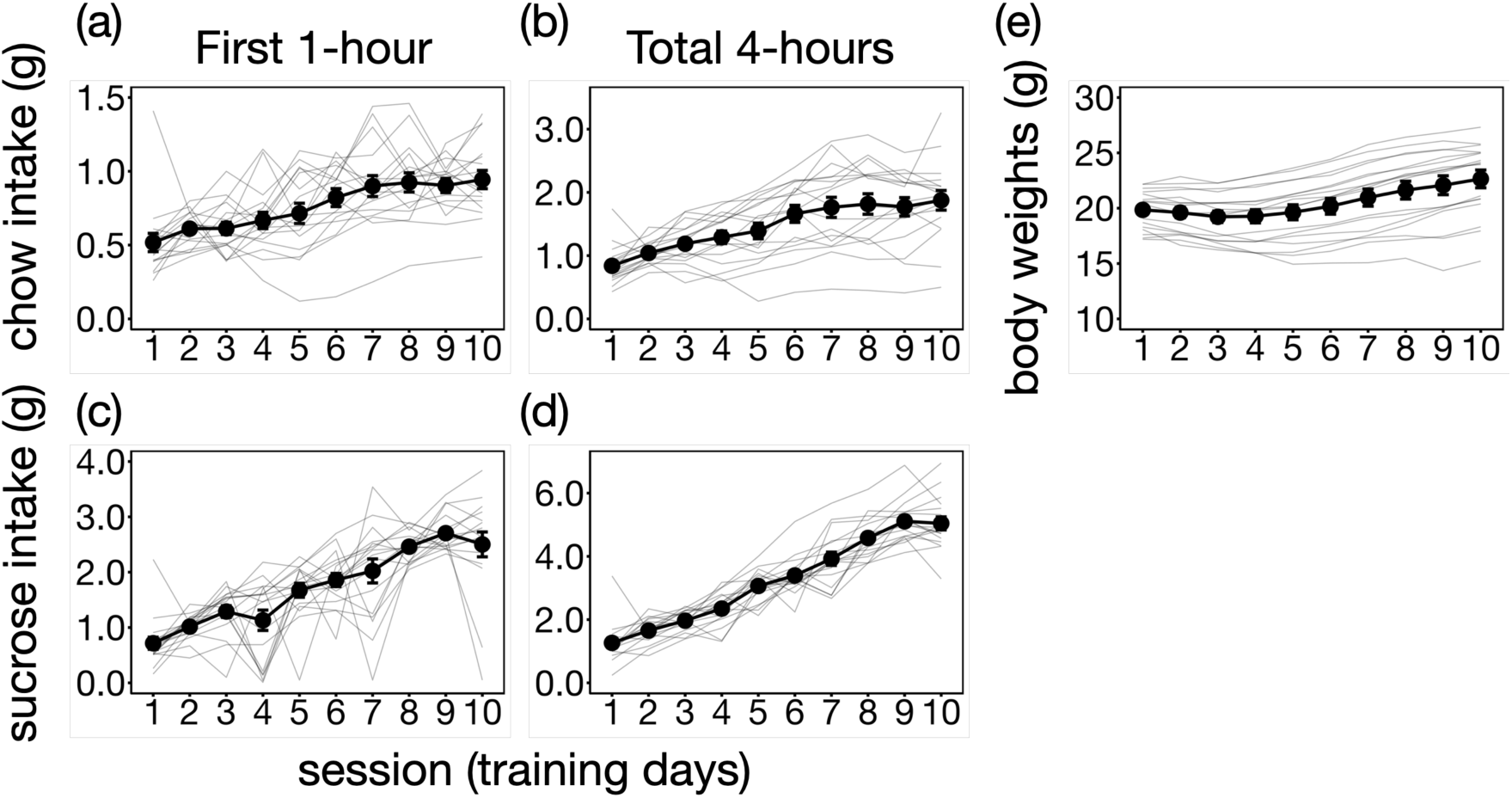
The average food consumption increased as the session proceeded and reached asymptotic levels. The thin gray lines represent individual data. The error bars denote standard errors. (a) 1-hour consumption of normal chow. (b) Total consumption of normal chow up to 4-hour daily feeding sessions. (c) 1-hour consumption of sucrose solution given as palatable substance. (d) The 4 hours total consumption of sucrose solution. (e) The changes of body weights throughout the training sessions.

In the next day after accomplishing restricted feeding training (i.e., day-11), animals were tested the following procedure: at first, the animals were given 2-hours free access to the sucrose solutions to satiate (i.e., sensory-specific devaluation) without the context. The satiation has been commonly used in experiments of habit to judge a target instrumental behavior is sensitive to the values of an outcome (Dickinson & Balleine, 1994). Since satiation was conducted without context, a within-subject two-way ANOVA was performed on sucrose consumption during the first and second 1-hour periods on day 10 and the test day to examine whether consumption during satiation remained consistent (Figure 3a). Two mice that consumed less sucrose during the first 1 h of training were excluded from this analysis. These animals formed the feeding pattern that they consumed almost exclusively chows during the first 1-hour, but the satiation undertook only with the presentation of sucrose solution. The ANOVA revealed the significant interaction (F(1,14) = 27.92, p<.0001). Thus, t-tests were applied separately to the day-10 training and the test. These t-tests revealed the significant difference between the initial 1-hour and second 1-hour in both session (the day-10: t(14) = 10.20, p-value < .0001; the test: t(14) = 5.88, p <.0001).

**Figure 3.**
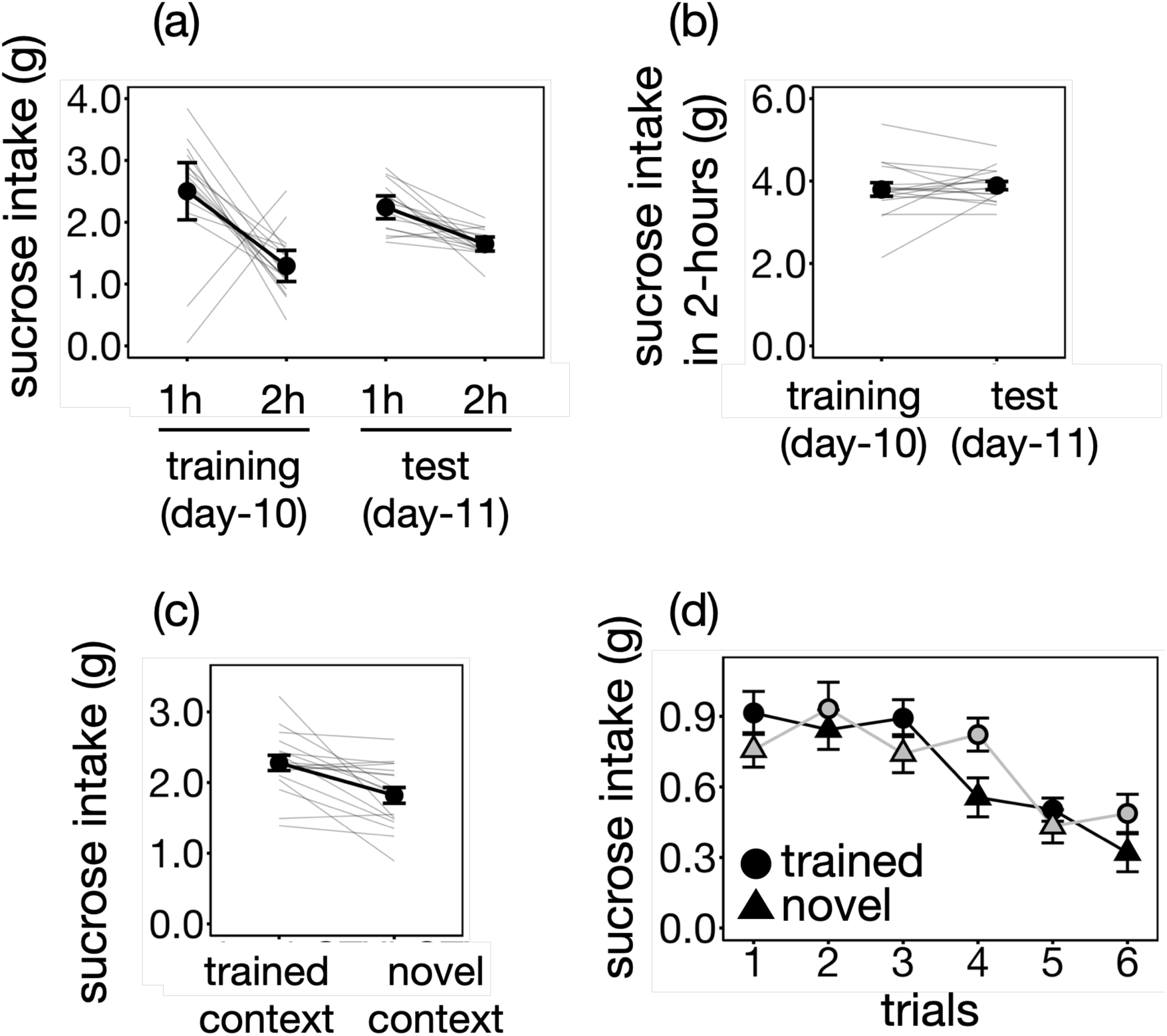
The contextually-induced binge-like consumption is resistant to sensory-specific devaluation by satiation of sucrose solution. (a) The consumption of sucrose solution during the satiation on the initial day 2 hours in day-10 and the test day. (b) the total sucrose solution intake over the course of two hours during the satiation. (c) the consumption of sucrose solution between the trained and novel context. (d) The trial-by-trial fluctuation of sucrose solution consumption under the trained and novel contexts revealed the context-specific elicitation of feeding behavior in the sated animals. The circles and triangles represent the trained and novel contexts, respectively. The black marker and line represent the group that began with the trained context, while the gray markers and line indicate the group that started with the novel context. The error bars represent SEM.

However, the slope of the decrease in consumption between the first and second 1-hour periods was shallower during the test, suggesting an influence of environmental changes (i.e., the absence of chow and context; Figure 3a). This may indicate a potential flaw in the satiation procedure, as it may not have sufficiently reduced hedonic needs for sucrose. Therefore, we conducted an additional t-test to compare total sucrose solution intake over the two-hour period between training day 10 and the satiation phase on the test day (Figure 3b). It revealed that no significant difference was observed between the days (t(16) = 0.63, p-value = 0.54). Therefore, total sucrose solution intake over the two-hour period was nearly identical between training and test sessions, suggesting that satiation was adequately achieved.

After the 2-hours free access to the sucrose solution, we asked whether binge-like overconsumption is resistant to devaluation of the palatable sugar. For that purpose, after the satiation, the animals were given access to sucrose solutions under the trained context or novel context. Each access continued 20 minutes, and the access was given in another context (trained-> novel or novel -> trained). Three alternations of context conditions (trained or novel) were conducted, resulting in a total of six trials and 1 hour eating occasion in the respective contexts. The total amounts of consumption given contexts were statistically significant (Figure 3c, right side, t(16) = 3.73, p = .002).Thus, the consumption was heightened in the trained context than the novel context. This effect can be observed that feeding behavior is influenced by specific contexts in the trial-by-trial fluctuation of consumption (Figure 3d). The linear mixed model revealed that both the trial and context was significant (χ²(5) = 76.40, p < .0001 for trials and χ²(1) = 12.32, p= .0004), while the significant interaction was not observed (χ²(5) = 2.01, p = .85).

Taken together, these results indicate that contextual enhancement can lead to the overconsumption of palatable food, even after 2 hours of satiation. However, it is important to consider whether this observation is specific to the binge-like behavior model. Since animals were trained with contextual appetitive conditioning over 10-days, with consistent pairing of context and food, the context itself might acquire a controlling effect that elicits feeding behavior beyond hedonic needs. In other words, mere contextual conditioning might be sufficient to influence excessive food intake. Experiment 2 was designed as a control experiment to investigate this possibility.

### Experiment 2: specificity of contextual control in the binge-eating model

The sucrose and chow consumption were developed across days (Figure 4). A one-way analysis of variance (ANOVA) revealed the significant effect of 1-hour consumption of sucrose solution (F(3.53,56.43) = 36.87, p < .0001), that of total 4-hours (F(2.47,39.54) = 40.37, p < .0001), 1-hour consumption of chow (F(9.0,144.0) = 11.41, p < .0001), and that of total 4-hours (F(9.0,144.0) = 13.30, p < .0001). Even though chow was ad libitum in the experiment 2, both chow and sucrose consumption were significantly increased as the session proceeded.

**Figure 4.**
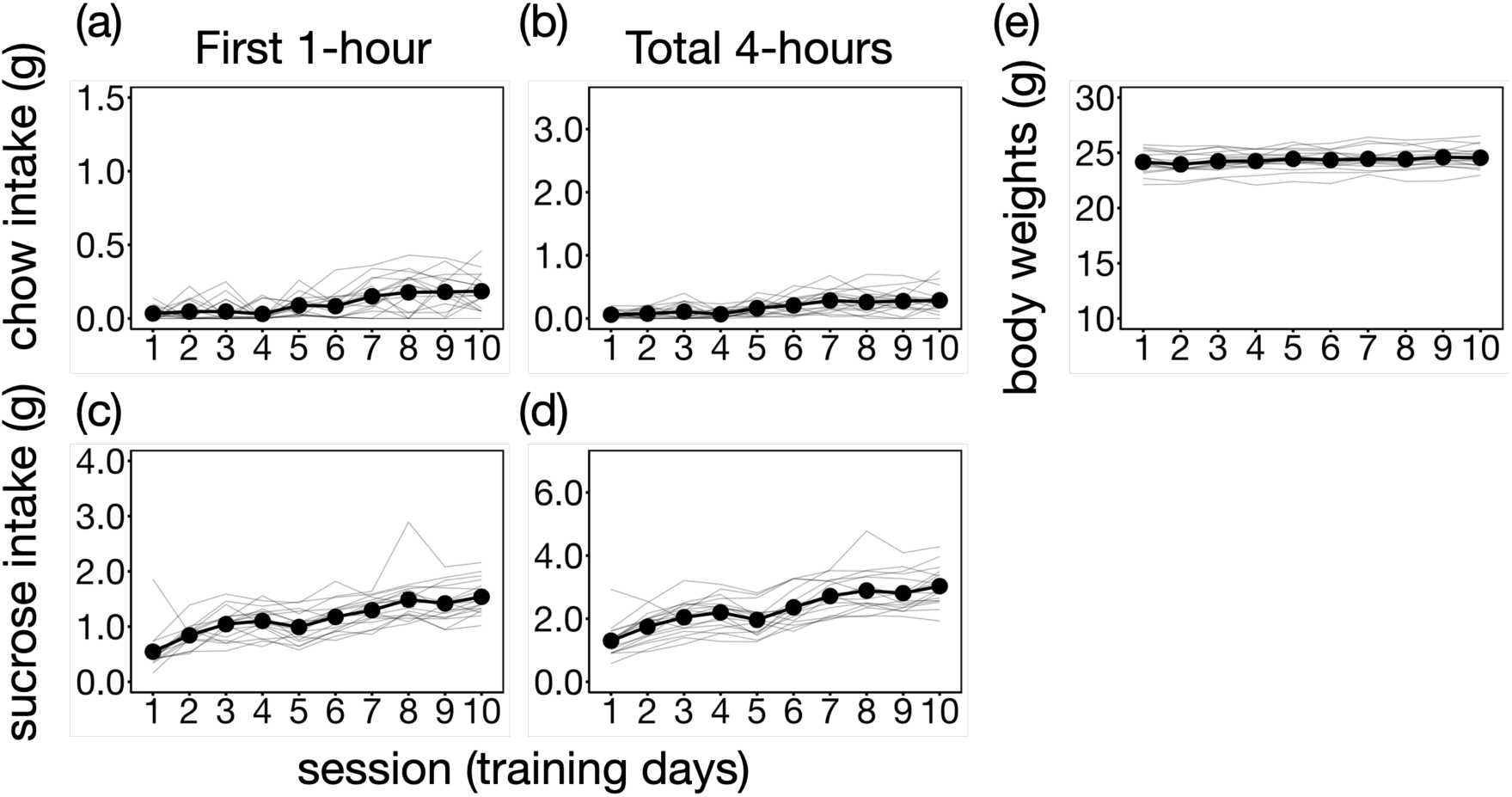
The average food intake remained at a lower level compared to Experiment 1. The thin gray lines represent individual data. The error bars denote standard errors. The ranges of the vertical axis were set the same as Figure 2. (a) 1-hour consumption of normal chow. (b) Total consumption of normal chow up to 4-hours daily feeding sessions. (c) 1-hour consumption of sucrose solution given as palatable food. (d) The 4 hours total consumption of sucrose solution. (d) The 4 hours total consumption of sucrose solution. (e) The changes of body weights throughout the training sessions.

However, when food consumption between experiment 1 and 2 was compared using two-way ANOVA (session × experiment), all measures in Figure 2 and 4 showed significant difference between experiment and interaction: 1-hour consumption of sucrose solution (experiment: F(1.00,32.00) = 11.67, p = .002, the interaction: F(4.60,147.08) = 6.43, p < .0001), that of total 4-hours (experiment: F(1.00,32.00) = 6.05, p = .02, the interaction: F(3.77,120.56) = 22.71, p < .0001), 1-hour consumption of chow (experiment: F(1.0,32.0) = 208.56, p < .0001, the interaction: F(5.33,170.53) = 3.03, p = .01), and that of total 4-hours (experiment: F(1.0,32.0) = 127.27, p < .0001, the interaction: F(3.43,109.67) = 11.70, p < .0001). These results demonstrate that the amount of food consumption grew up more in the restricted feeding than under the free access to the chow. Thus, binge-like food consumption was notable in Experiment 1, but not in Experiment 2. The body weights significantly changed across training sessions (F(3.44,55.07) = 6.46, p < .0005). However, the multiple comparison revealed there was no systematic differences (session 2 was lower than 3, 5, 9, and 10, p-values ranged from 0.003 to 0.018).

The test was conducted the next day after 10 days training, and the testing procedure was completely the same as experiment 1. Thus, the same analysis was applied to the results of experiment 2. The 2-hours of satiation decreased the sucrose consumption similarly to that of the training (Figure 5a). The two-way ANOVA revealed only the first and second hours were significant (F(1,16) = 215.80, p < .0001), but not the days (day-10 and test: F(1,16) = 2.76, p =.12) and the interaction (F(1,16) = 0.81, p = .38). The total consumption during two hours also did not show the significant difference between the training day-10 and the satiation periods in the test (t(16) = 1.66, p = .11), revealing that the satiation effectively reduced the sucrose consumption (Figure 5b).

**Figure 5.**
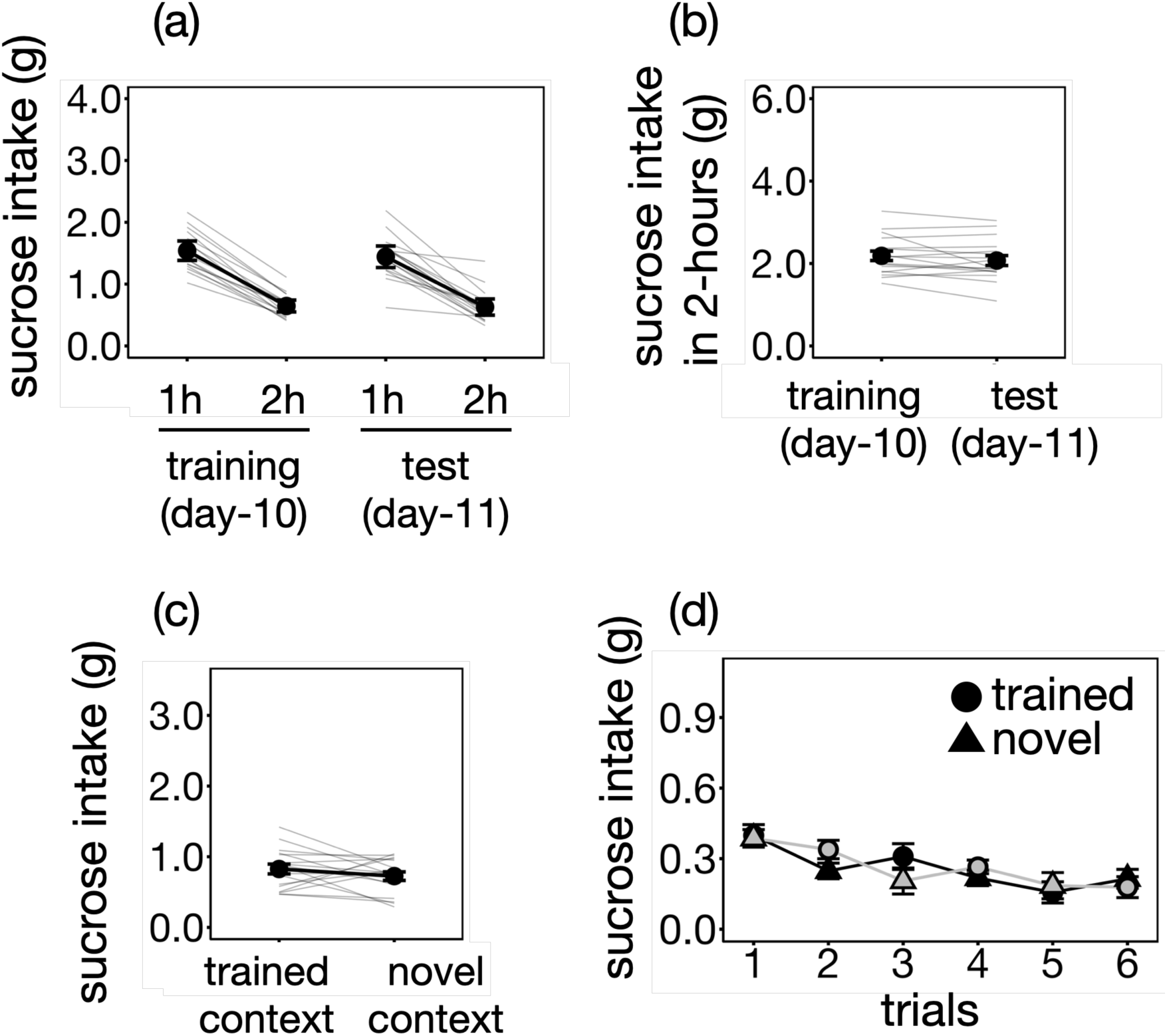
The mere contextual conditioning did not facilitate feeding behavior resistant to the satiation. The ranges of the vertical axis were set the same as Figure 3. The contextually-induced binge-like consumption is resistant to sensory-specific devaluation by satiation of sucrose solution. (a) The consumption of sucrose solution during the satiation on the initial day 2 hours in day-10 and the test day. (b) the total sucrose solution intake over the course of two hours during the satiation. (c) the consumption of sucrose solution between the trained and novel context. (d) The trial-by-trial fluctuation of sucrose solution consumption under the trained and novel contexts revealed the context-specific elicitation of feeding behavior in the sated animals. The circles and triangles represent the trained and novel contexts, respectively. The black marker and line represent the group that began with the trained context, while the gray markers and line indicate the group that started with the novel context. The error bars represent SEM.

Unlike Experiment 1, the sucrose consumption under the trained or novel context did not produce the significant difference (the right side of Figure 5c; t(16) = 1.21, p = .24). Similarly, the trial-by-trial fluctuation also demonstrated that the sucrose consumption did not differ between the trained and novel context (Figure 5d). Indeed, the linear mixed model revealed that only the trial was significant (χ²(5) = 39.47, p < .0001), while the significant effect of the context was not observed (χ²(1) = 1.93, p = .16). The interaction between these neither significant (χ²(5) = 5.57, p = .35). These results indicated that the contextual enhancement was not found in the animals that did not develop the binge-like consumption of the palatable food.

*Experiment 3*: the mice did not exhibit context-specific sucrose consumption in Experiment 2, suggesting that the binge-like drinking protocol is necessary for this effect. However, an alternative interpretation is that Pavlovian learning did not occur because the animals were not food-deprived, resulting in a failure to form the association. To examine this possibility, Experiment 3 employed a CPP task. Specifically, the animals’ preference between the trained and novel contexts was assessed both before and after experiencing sucrose-context pairing. If mice successfully formed an association, they would exhibit a greater preference for the trained context even in the absence of food deprivation.

The training procedure was completely the same as that of Experiment 2. The food consumption during 10-days training is similar (Figure 6). A one-way ANOVA revealed the significant effect of 1-hour consumption of sucrose solution (F(9,72) = 23.27, p < .0001), that of total 4-hours (F(2.47,39.54) = 12.40, p < .0001), 1-hour consumption of chow (F(9.0,72.0) = 3.18, p = .003), and that of total 4-hours (F(9.0,72.0) = 4.809, p < .0001). Again, although food was ad libitum, the intakes slightly increased across the session.

**Figure 6.**
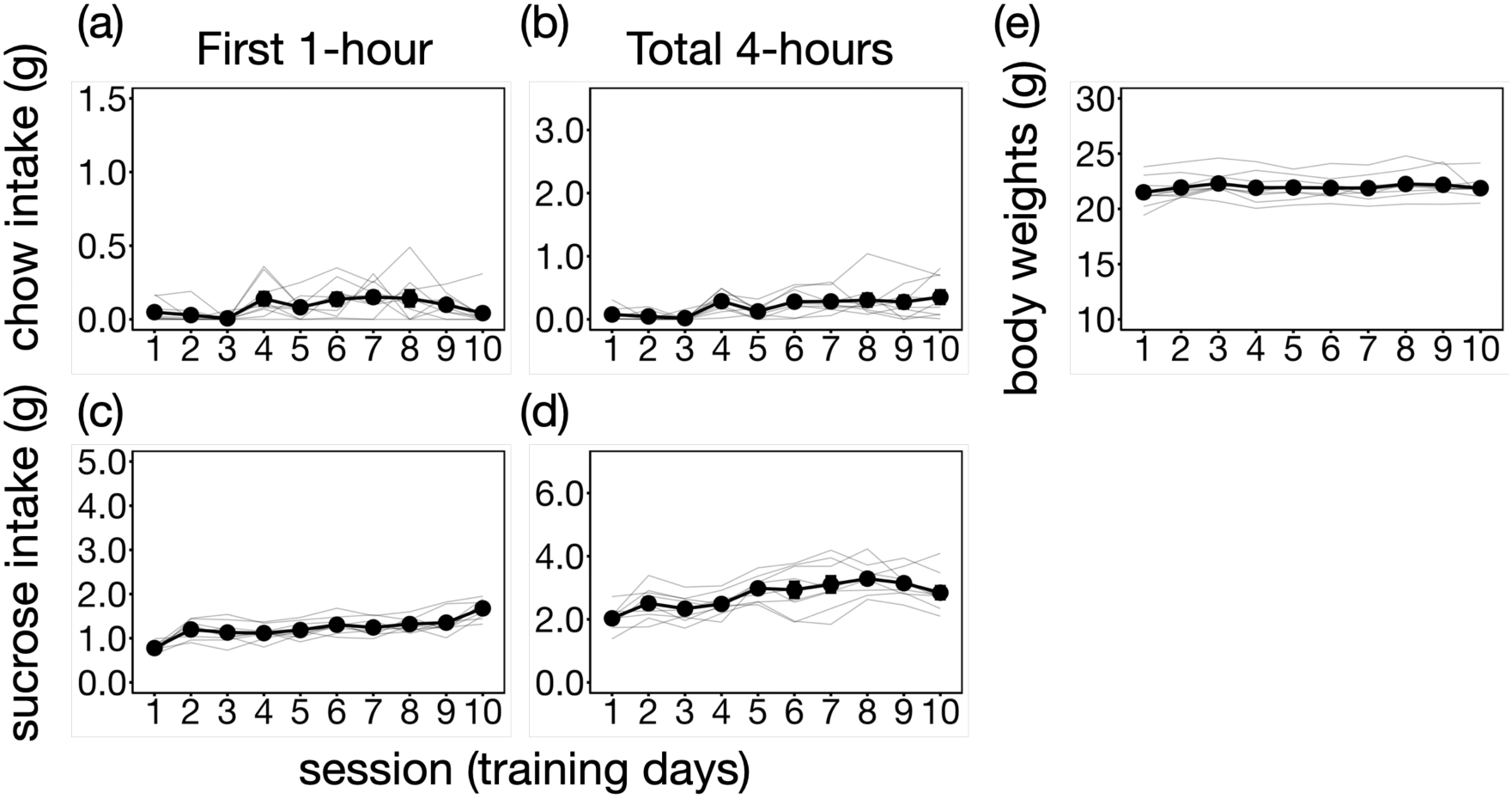
The chow and sucrose intakes in Experiment 3. The thin gray lines represent individual data. The error bars denote standard errors. The ranges of the vertical axis were set the same as Figure 2. (a) 1-hour consumption of normal chow. (b) Total consumption of normal chow up to 4-hours daily feeding sessions. (c) 1-hour consumption of sucrose solution given as palatable food. (d) The 4 hours total consumption of sucrose solution. (d) The 4 hours total consumption of sucrose solution. (e) The changes of body weights throughout the training sessions.

The place preference between contexts were examined to test whether Pavlovian learning was established during training (Figure 7). The t-test revealed the significant difference between pre- and post-tests (t(8) = 2.63, p = .03). Therefore, it was likely that the animals learned the positive association regarding the context even in the absence of physiological motivation during the course of training.

**Figure 7.**
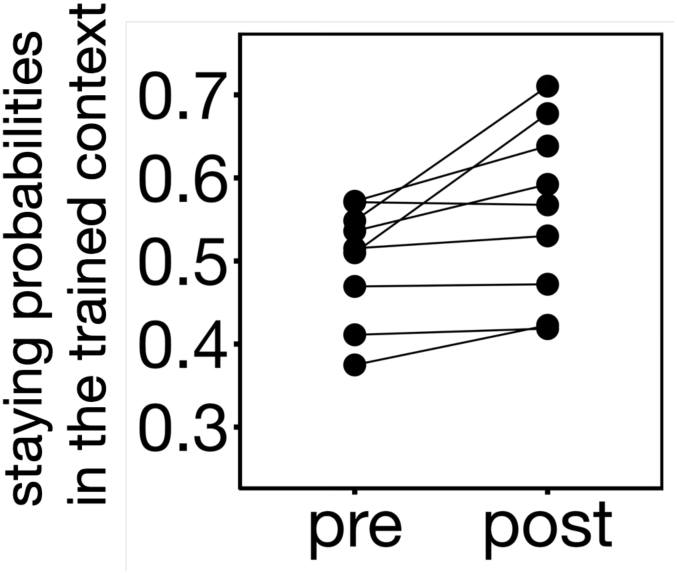
The staying probabilities in the trained context was compared between before (“pre” at day-0) and after training (“post” at day-11 test).

**Figure 8.**
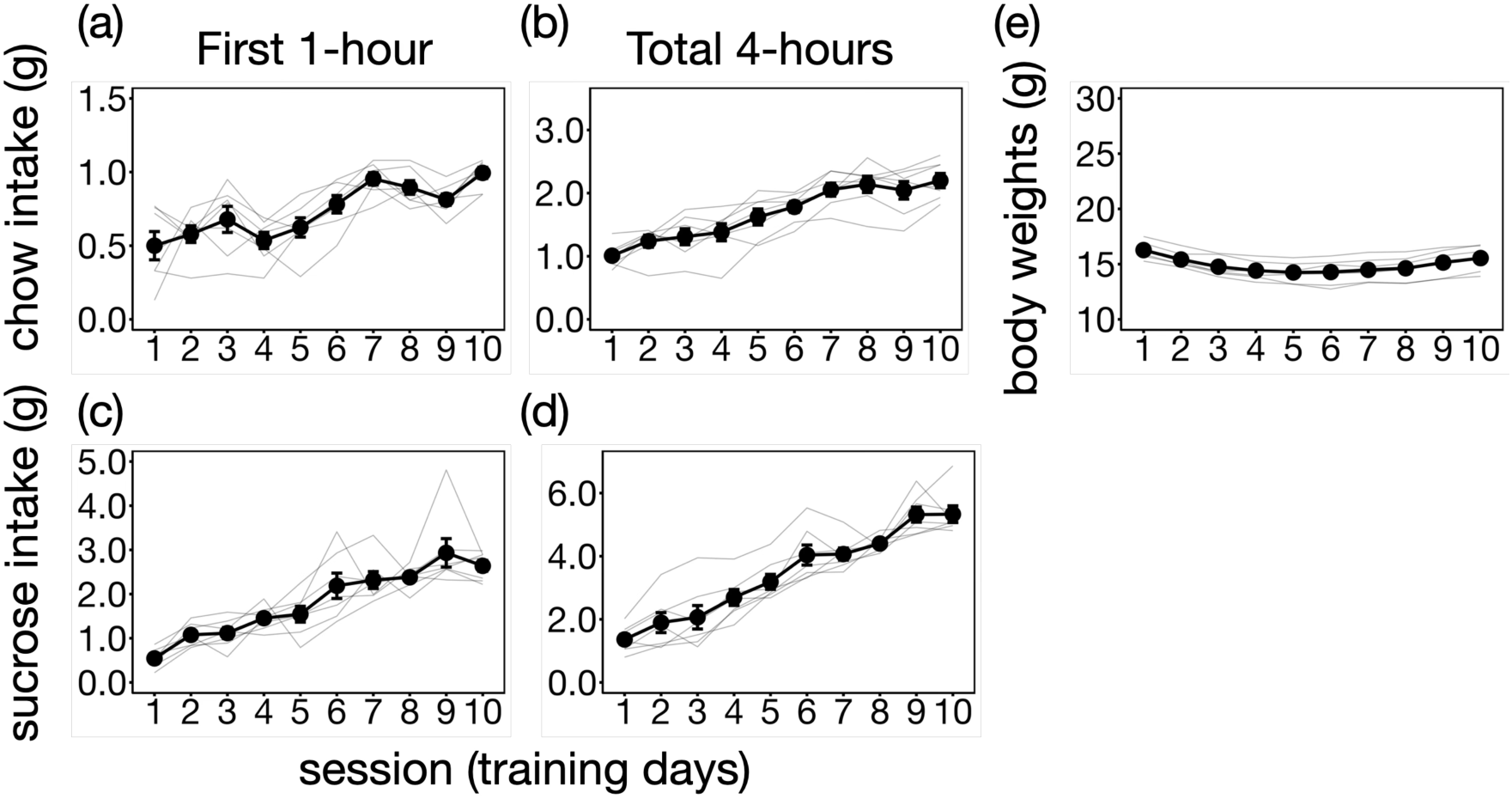
The average food consumption increased as the session proceeded and reached asymptotic levels. The thin gray lines represent individual data. The error bars denote standard errors. (a) 1-hour consumption of normal chow. (b) Total consumption of normal chow up to 4-hour daily feeding sessions. (c) 1-hour consumption of sucrose solution given as palatable substance. (d) The 4 hours total consumption of sucrose solution. (e) The changes of body weights throughout the training sessions.

*Experiment 4*: a critical limitation of the experiments described above is that only male mice were used, potentially introducing bias in the results. Additionally, the experiments were susceptible to the confounding effect of neophobia: the animals may have been simply reluctant to consume sucrose (Experiments 1 and 2) or to remain in the novel context (Experiment 2) due to unfamiliarity rather than learned associations. To address these issues, female mice were trained using a modified procedure in which they were exposed to one positive trained context (CTX+) during feeding and another negative context (CTX −) after the feeding session.

One animal did not increase food intake and excluded from experiment to avoid starvation. A one-way ANOVA revealed the significant effect of 1-hour consumption of sucrose solution (F(9.0,54.0) = 24.18, p < .0001), that of total 4-hours (F(9.0,54.0) = 67.38, p < .0001), 1-hour consumption of chow (F(9.0,54.0) = 10.87, p < .0001), and that of total 4-hours (F(9.0,54.0) = 37.94, p < .0001). Similar to Experiment 1 in male, the sucrose intake was tripled during the course of training in female.

The test was performed in the same manner as Experiment 1. Unlike Experiment 1, the 2-hour satiation did not reduce consumption (Figure 9a). The two-way ANOVA revealed a significant interaction between sessions and satiation times. (F(1,6) = 9.56, p=.021). The subsequent comparison showed the significant difference between sucrose intakes during 1- and 2-hours in the day-10 training (t(6) = 5.34, p = .002) but not in the satiation session in the test (t(6) = .57, p = .59). It resulted in heightened sucrose intakes during 2-hours between the last training and test sessions (t(6) = 11.73, p < .0001; Figure 9b). This result was completely unexpected and may reflect sex differences in binge-like consumption (see General Discussion). However, we must remain agnostic as to whether the animals were sufficiently sated in the operational sense.

**Figure 9.**
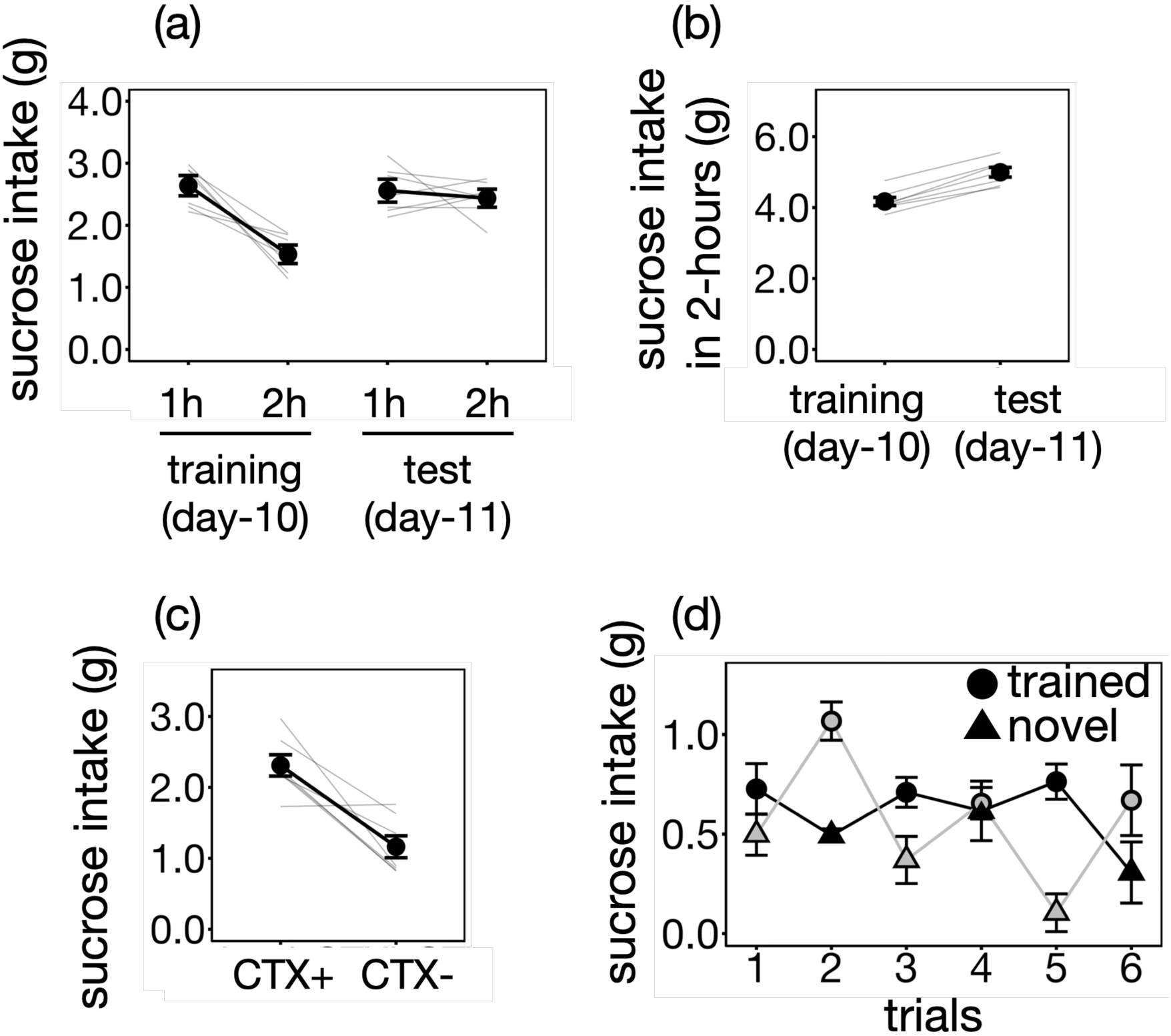
The replication with the modified procedure for Experiment 1 using female mice. (a) The consumption of sucrose solution during the satiation on the initial day 2 hours in day-10 and the test day. (b) the total sucrose solution intake over the course of two hours during the satiation. (c) the consumption of sucrose solution between the trained and novel context. (d) The trial-by-trial fluctuation of sucrose solution consumption under the trained and novel contexts revealed the context-specific elicitation of feeding behavior in the sated animals. The circles and triangles represent the trained and novel contexts, respectively. The black marker and line represent the group that began with the trained context, while the gray markers and line indicate the group that started with the novel context. The error bars represent SEM.

Nevertheless, the context-specific enhancement of sucrose consumption remained notable in the subsequent tests (Figure 9cd). The total sucrose consumption was higher in the CTX+ context compared to the CTX-(t(6) = 4.66, p = .003), indicating that contextual control effectively modulated sucrose intake, similar to Experiment 1. The linear mixed model revealed significant session effects on the trial-by-trial fluctuation in sucrose consumption. (χ²(5) = 12.70, p = .03) and the contexts (χ²(1) = 29.12, p < .0001).

These results replicate the key finding of Experiment 1: the trained context enhanced sucrose consumption after 2 hours of satiation. The modifications in Experiment 4 compared to Experiment 1 were (1) the use of female mice and (2) the exposure to a non-reinforced context instead of a novel one. Thus, the results collectively support the generality of contextual control across sexes and rule out the possibility that the effect was confounded by neophobia, which could have hindered the animals’ sucrose consumption in the novel context.

However, two notable differences were observed between Experiment 1 and Experiment 4. First, female mice did not decrease sucrose consumption during the 2-hour satiation session (Figure 9a), in contrast to male mice (Figure 3a). Another difference was that the context continued to facilitate consumption throughout the test trials in female mice (Figure 9d). In male mice, sucrose consumption under the trained context decreased by approximately half by the end of the trials (fifth or sixth), compared to the initial trials (first or second). Despite the significant session effect, female mice exhibited only a moderate decrement in consumption across trials, indicating persistent drinking. Since females have smaller body weights than males, absolute body weight does not explain this sex difference.

## General Discussion

The present study aimed to determine whether binge-like consumption in a mouse model is subject to contextual enhancement of palatable sugar intake and whether feeding behavior beyond satiation depends on the context in which binge eating is developed. To investigate these possibilities, we employed a restricted feeding protocol to induce overconsumption within a specific context and subsequently assessed whether the mice would seek sucrose solutions even after a 2-hour period of satiation.

Experiment 1 demonstrated that animals were more likely to consume sucrose solutions in the context where they had developed binge-like overconsumption. This contextual enhancement was specific to the context associated with binge eating (Figure 3c-d), as the novel context did not exert the same regulatory influence. Since sucrose consumption decreased over the course of satiation, it suggests that the animals had access to an adequate amount of palatable sugar (Figure 3a). Thus, exposure to the trained context facilitated food consumption even after satiation. This context-specific facilitation of overconsumption was not observed in Experiment 2, where animals had free access to standard chow during the training, resulting in the absence of binge-like eating.

However, the mice in Experiment 2 were not food-deprived, which may have led to a lack of motivation to form a Pavlovian association. In fact, food deprivation has been reported to facilitate the formation of Pavlovian conditioning and enhance the discriminability of conditioned stimuli (Chu & Ågmo, 2012; Tabbara et al., 2016).

Nevertheless, another study demonstrated successful conditioning in animals without food restriction (Weingarten, 1983). In the present study, exposure to the context did not evoke feeding behavior during the sucrose intake test. However, when location preference was measured in Experiment 3, the mice increased their time spent in the trained context. This result suggests that, even in the absence of food motivation, a learning signature related to the context was still observed. Therefore, it is unlikely that the lack of motivation in Experiment 2 accounted for the results.

Our Experiment 1 had two limitations. First, only male mice were used. Given that binge-eating disorder is more prevalent in females—not only due to socio-cultural factors (Davis, Graham & Wilders, 2020) but also physiological ones (Asarian & Geary, 2013; Novelle & Diéguez, 2019)—examining sex differences is crucial. The second issue was neophobia. Because sugar consumption was compared between a trained and a novel context, the latter might have suppressed eating rather than the former facilitating it. To address these issues, Experiment 4 was conducted. The results largely mirrored those of Experiment 1 but also revealed a sex difference. In contrast to Experiment 1, two hours of satiation did not reduce sugar solution consumption (Figure 9a). Indeed, this extended drinking period resulted in a greater intake compared to the final day of training (Figure 9b), raising uncertainty as to whether the mice were truly sated. Nevertheless, contextual control over sugar intake remained evident and was even more pronounced in female mice than in males. The rodent model of binge-eating also revealed female rats are more susceptible than males (Klump, Racine, Hildebrandt & Sisk, 2013). However, in ther present study, obsered larger effect may have stemmed from differences in testing procedures: Experiment 1 used a novel context, whereas Experiment 4 employed a negative context (CTX-). Therefore, it is most prudent to conclude that contextual control over sugar intake was observed in both sexes.

The observed lack of sensitivity to the reinforcer resembles habitual behavior in instrumental learning (Dickinson et al., 1983). It is well-established that behaviors developed through both classical and instrumental learning are typically sensitive to the value of the reinforcer (Adams & Dickinson, 1981; Holland, 1981; Rescorla, 1973). The outcome devaluation, including satiation or taste aversion learning, are commonly used to reduce the value of the reinforcer, which is known to diminish the animal’s response (Colwill & Rescorla, 1985). This mode of behavior is termed ‘goal-directed’ because it is motivated by the value of the outcome to fulfill specific needs (Dickinson & Balleine, 1994). In contrast, habitual behavior emerges through extended learning periods, and is characterized by insensitivity to the value of the outcome (Dickinson, 1985). For instance, instrumental responses can persist even after the reinforcer has been devalued (Dickinson et al., 1983). This form of habitual behavior has also been observed in human studies (Cushman & Morris, 2015; Tricomi, Balleine, & O’Doherty, 2009). Additionally, behavioral automaticity resulting from extended training has been documented in Pavlovian learning in crickets (Mizunami et al., 2019). In these studies, behaviors were considered automated and insensitive to the value of the behavioral consequence due to repeated exposure to specific environment-behavior cycles, similar to the contextually-induced feeding observed in our study. This view of value-insensitivity is supported by a theoretical model that describes the development of habitual behavior (Yamada & Toda, 2023).

The physiological mechanisms underlying habitual behavior involve the dorsolateral striatum, which is crucial for generating automated behaviors, whereas the dorsomedial striatum is more associated with goal-directed behavior (Gremel & Costa, 2013; Yin, Knowlton & Balleine, 2004). The dorsolateral striatum is also implicated in binge eating behavior (Furlong et al., 2014). Abnormalities in striatal function have been reported in patients with binge-eating disorder (Kessler et al., 2016; Wang et al., 2011). Striatal circuits involved in behavioral automaticity overlap with those implicated in other disorders, such as obsessive-compulsive disorder and drug addiction (Kenny, 2011; Simmler & Ozawa, 2019). For instance, striato-midbrain-striatal connectivity plays a critical role in habitual cocaine-seeking behavior (Belin & Everitt, 2008). These circuits may serve as a common pathway for executing automated behaviors driven by environmental cues, including the overconsumption elicited by contextual factors observed in our study.

The functional impairments specific to binge eating may reside in the mechanisms governing feeding regulation. Normally, feeding is terminated by detecting stimulation of the gut (Aitken et al., 2024; Kim et al., 2024). In binge eating, however, this feeding termination may be disrupted by other regulatory mechanisms that promote food intake. Recent research, indeed, has shown that exposure to food-related contexts can enhance appetite through increased activity in the lateral hypothalamus (Subramanian et al., 2023).

Additionally, sympathetic nerve activity has been found to inhibit the release of glucagon-like peptides 1, further promoting appetite (Ren et al., 2024). It is possible that these top-down processes, rather than bottom-up signals from the gut, likely involve context-specific amplification of eating behaviors. However, the mechanisms by which contextual factors drive food consumption beyond satiety remain poorly understood. Further studies are needed to elucidate how environmental factors modulate feeding regulation.

As discussed above, while there may be certain commonalities between overeating and habits, it is important to consider the methodological differences; in studies of habitual instrumental behavior, animals typically undergo extended training followed by sensory-specific devaluation, with an immediate extinction test (e.g., Adams & Dickinson, 1981; Thrailkill & Bouton, 2015). In the extinction test, the animals do not experience the outcome event (e.g., sucrose solution or pellet) while responding, thus not having the opportunity to update the learning about the association between response and outcome. This is a notable methodological difference compared to our experiment, where sucrose was continuously presented and mice could freely consume it during the test. In our experiment, the mice had the opportunity to monitor their needs and adjust their behavior each time feeding behavior occurred. Nevertheless, what actually happened was that, through repeated trials, the mice showed increased sucrose intake in the trained context. Therefore, the automaticity of the behavior observed in our study is unlikely to be an artifact of procedural differences, as the apparent effect. However, some issues remain to be considered. While the dorsal striatum is a common neural substrate involved in both habits and overeating, the procedural and phenomenological differences between experiments may reflect differences in upstream mechanisms, which influence behavior closer to the striatum. Future studies on binge eating should investigate how contextual stimuli influence the mechanisms of sucrose craving before the behavior occurs, and reveal the neural processes that is specifically malfunctioning in binge eating disorder.

Lastly, our study has certain limitations. In the training procedure devised by Yasoshima & Shimura (2015), binge eating was characterized by the consumption of sucrose solution in amounts exceeding the expected water intake based on the amount of food consumed. To support this, they included a group that was presented with chow and water only between 9:00 and 13:00, allowing a comparison of sucrose and water intake. In our study, since we did not include this group, it is important to note that the observed increase in sucrose intake is comparable to that in previous studies. Additionally, in our experiments, chow and sucrose were always presented together with the context, while only sucrose was used in the test phase. As a result, although feeding behavior was induced by the trained context, it remains unclear whether this effect was specifically tied to the palatable sucrose. Future research should investigate the extent to which contextual control of behavior generalizes across different food types.

In summary, the present experiments demonstrated that the context in which binge-like behavior developed facilitated feeding beyond satiety. This effect was context-specific and resembled habitual behavior in instrumental learning. However, simply establishing a Pavlovian association between the context and palatable substance was insufficient for eliciting contextual facilitation of overconsumption in our study. Instead, contextual facilitation may emerge when food restriction is combined to binge-like behavioral abnormalities.

## Acknowledgements

The present research was funded by JSPS KAKENHI (24K16867 to H.M.).

## Declaration of generative AI

The authors used ChatGPT 4.0 for language-proofing to refine the manuscript written by non-native English speakers. After using this tool/service, the authors reviewed and edited the content as needed and takes full responsibility for the content of the publication.

